# Gene model for the ortholog of *Ilp4* in *Drosophila grimshawi*

**DOI:** 10.1101/2025.08.25.672213

**Authors:** Rachael Cowan, Alyssa Koehler, Clairine I. S. Larsen, Jeffrey S. Thompson, Andrew M Arsham, Lori Boies

## Abstract

Gene Model for the *Insulin-like peptide 4 (Ilp4)* ortholog in the *D. grimshawi (dgriCAF1*) assembly (GeneBank Accession: GCA_000005155.1) of *Drosophila grimshawi*. The characterization of this ortholog was conducted as part of a broader, developing dataset aimed at investigating the evolution of the Insulin/insulin-like growth factor signaling (IIS) pathway across the genus *Drosophila*, using the Genomics Education Partnership gene annotation protocol within Course-based Undergraduate Research Experiences.

## Introduction

> *This article reports a predicted gene model generated by undergraduate work using a structured gene model annotation protocol defined by the Genomics Education Partnership (GEP; thegep.org) for Course-based Undergraduate Research Experience (CURE). The following information in quotes may be repeated in other articles submitted by participants using the same GEP CURE protocol for annotating Drosophila species orthologs of Drosophila melanogaster genes in the insulin signaling pathway*.

The insulin signaling pathway is well characterized in *Drosophila* and is highly conserved across metazoans. Owing to this strong evolutionary conservation, *Drosophila* serves as a valuable model for studying insulin signaling in humans (Biglou, *et. al*. 2021). The *insulin-like peptide 4 (Ilp4)* gene has been shown to have high expression within the mesoderm of the embryo and the midgut in the larval stage (Brogiolo, *et. al*., 2001). Of the eight insulin-like peptides within the *Drosophila* insulin signaling pathway, *Ilp4 1 - 7* are highly conserved across species, with *Ilp4 and Ilp7* being the most conserved (Grönke, *et. al*., 2010). While highly conserved, there is scant evidence of the specific function of *Ilp4* within the scientific literature. The model presented here is the ortholog of *Ilp4* in the dgriCAF1 assembly of *D. grimshawi* (GCA_000005155.1 - Clark et al. 2007) and corresponds to the XP_032590595.1 predicted model *D. grimshawi* (LOC6556940). This gene model is based on RNA-seq data from *D. grimshaw*i (SRP073087) in *D. melanogaster* from *FB2022_02* (Larkin *et al*., 2021).

*Drosophila grimshawi* (NCBI:txid7222) belongs to the Picture Wing group of Hawaiian *Drosophila*, as defined by Kaneshiro and colleagues (1995). Phylogenetic studies indicate that the Hawaiian *Drosophila* form a monophyletic lineage that is sister to the *Scaptomyza* clade, with both lineages situated within the *Drosophila* subgenus of the genus *Drosophila* (Kambysellis et al., 1995; Baker & DeSalle, 1999). The group derives its name from the striking pigmentation patterns on their wings. *D. grimshawi*, first described by Oldenberg in 1914, inhabits high-elevation, cool tropical rainforests on the Maui Complex islands, where it reproduces on decaying plant material (Carson et al., 1970; Carson, 1983)

“In this GEP CURE protocol students use web-based tools to manually annotate genes in non-model Drosophila species based on orthology to genes in the well-annotated model organism, the fruit fly Drosophila melanogaster. This allows undergraduates to participate in course-based research by generating manual annotations of genes in non-model species (Rele et al., 2023).

Computational-based gene predictions in any organism are often improved by careful manual annotation and curation, allowing for more accurate analyses of gene and genome evolution (Mudge and Harrow, 2016; Tello-Ruiz et al., 2019). These models of orthologous genes across species, such as the one presented here, then provide a reliable basis for further evolutionary genomic analyses when made available to the scientific community.” (Myers et al., 2024).

## Results

### Synteny

*Ilp4* occurs on *chromosome 3L* in *D. melanogaster* and is flanked by *I-2, CG43897* with *Ilp5* nested within, *CG32052* with *Ilp3* and *Ilp2* nested within, *Ilp4, Ilp1, Zasp67*, and *Ae2*. We determined that the putative ortholog of *Ilp4* is found on scaffold CH916366 in *D. grimshawi* at LOC6556940 (via *tblastn* search with an e-value of 8e-09 and percent identify of 37.04%), where it is surrounded by genes with the following locus IDs: *LOC6556854, LOC6556919* with *LOC6557001* nested within, *LOC6557056* with *LOC6556947, LOC6556893*, and *LOC6556922* nested within, *LOC6556876, LOC6556918*, and *LOC6556946*. which corresponds to *I-2, CG43897* with *Ilp5* nested within, *CG32052* with *Ilp3* and two *Ilp2* paralogs nested within, *Ilp1, Zasp67*, and *Ae2 in D. melanogaster* as determined by *blastp* (Figure 1A, Altschul et al., 1990). We believe this is the correct ortholog assignment for *Ilp4* in *D. grimshawi* because local synteny is conserved with the exception of there being a duplication event resulting in two paralogs of *Ilp2*.

**Figure 1:**
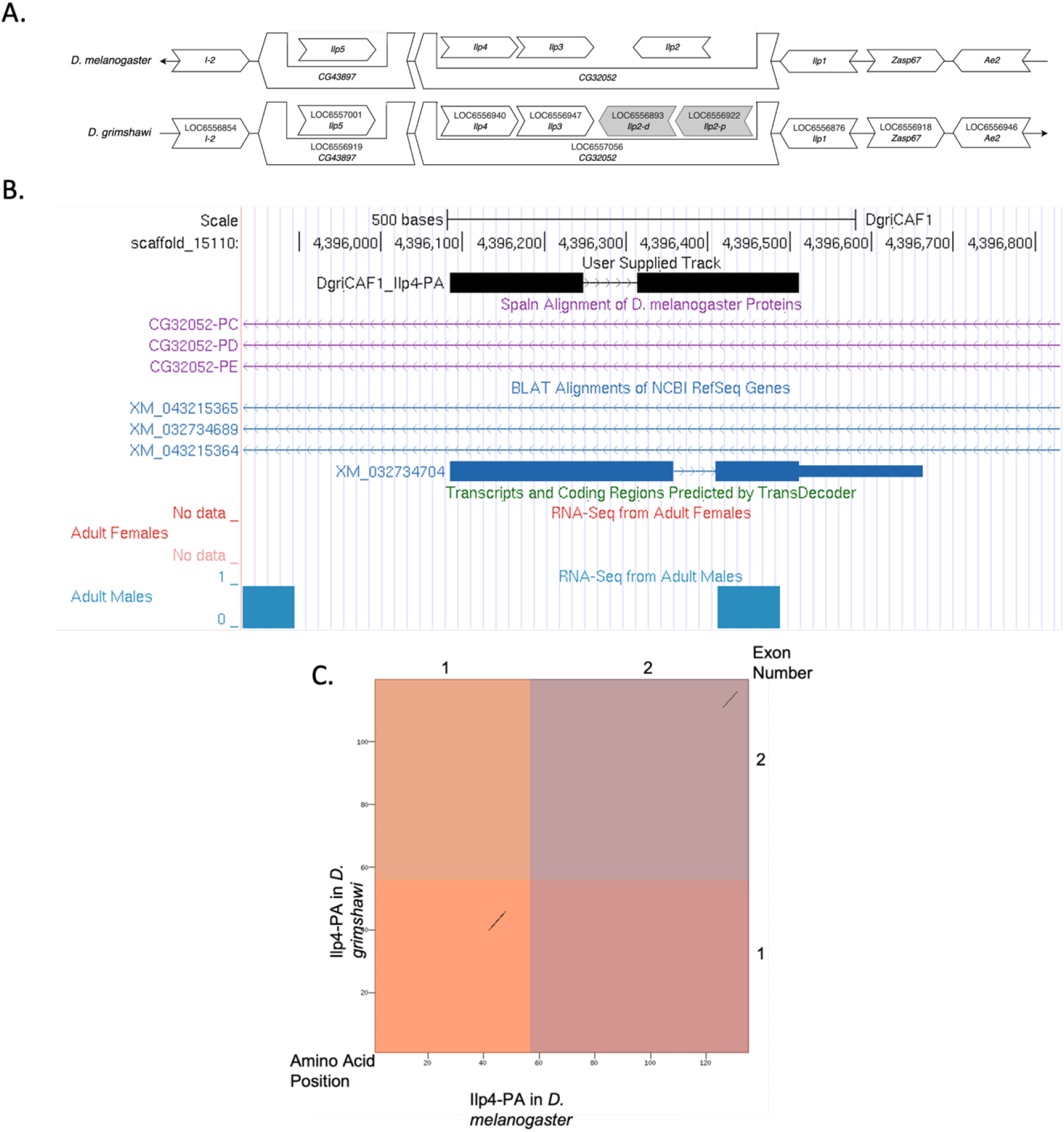
Ilp4 gene model comparison between Drosophila grimshawi and Drosophila melanogaster. **(A) Local synteny of genomic neighborhood of Ilp4 in D. melanogaster and D. grimshawi.** Gene arrows pointing in the same direction as the target gene *(Ilp4)* in both *D. grimshawi* and *D. melanogaster* are on the same strand as the target gene; gene arrows pointing in the opposite direction are on the opposite strand. The thin underlying arrows pointing to the right indicate that *Ilp4* is on the + strand; arrows pointing to the left indicate that *Ilp4* is on the – strand. White arrows in *D. grimshawi* indicate the locus ID and the orthology to the corresponding gene in *D. melanogaster*. Gray arrows indicate paralogs of a duplication of downstream gene *Ilp2* that is not present in *D. melanogaster*. The gene names given in the *D. gimshawi* gene arrows indicate the orthologous gene in *D. melanogaster*, while the locus identifiers are specific to *D. grimshawi*. **(B) Gene Model in UCSC Track Hub (Raney et al. 2014):** the gene model in *D. grimshawi* (black), Spaln of D. melanogaster Proteins (purple, alignment of refseq proteins from *D. melanogaster*), BLAT alignments of NCBI RefSeq Genes (blue, alignment of refseq genes for *D. grimshawi*), RNA-Seq from Adult males (blue) and adult females (red), alignment of Illumina RNAseq reads from *D. grimshawi*), and Transcripts (green) including coding regions predicted by TransDecoder and Splice Junctions Predicted by regtools using *D. grimshawi* RNA-Seq (SRP073087). The custom gene model (User Supplied Track) is indicated in black with exon depicted with wide boxes, intron with narrow lines (arrows indicate direction of transcription). **(C) Dot Plot of Ilp4-PA in *D. melanogaster* (*x*-axis) vs. the orthologous peptide in *D. grimshawi* (*y*-axis)**. Amino acid number is indicated along the left and bottom; exon number is indicated along the top and right, and exons are also highlighted with alternating colors. There are large gaps in the dot plot which indicates dissimilarity in the amino acid sequence between Ilp4-PA in *D. melanogaster* and *D. grimshawi*.

### Protein Model

*Ilp4* in *D. grimshawi* has 1 protein coding isoform (Ilp4-PA) (Figure 1B). Ilp4-PA contains 2 protein coding exons. *Ilp4* in *D. melanogaster* also has one isoform (Ilp4-PA) which also has two protein coding exons. The sequence of Ilp4-PA in *D. grimshawi* has 46.6% identity with the *Ilp4* in *D. melanogaster* as determined by *blastp* (Figure 1C). These data are also available in Extended Data files below, which are archived in CaltechData.

### Special characteristics of the protein model

#### Lack of Splice Junctions

There is no splice junction evidence track due to the relative lack of RNA-seq reads mapping to this region (Figure 1B). This may be because multiple *Ilp* paralogs with similar sequences impede accurate mapping of RNA-seq reads.

#### Low Sequence Similarity

There is low sequence similarity between *Ilp4* in *D. melanogaster* and *D. grimshawi* shown by *blastp* (46.6% identity), *tblastn* (37.04% identity) and the dot plot showing the protein alignment (Figure 1C). This is likely because *D. grimshawi* is highly diverged from *D. melanogaster*.

## Methods

“Detailed methods including algorithms, database versions, and citations for the complete annotation process can be found in Rele et al. (2023). Briefly, students use the GEP instance of the UCSC Genome Browser v.435 (https://gander.wustl.edu; Kent WJ et al., 2002; Navarro Gonzalez et al., 2021) to examine the genomic neighborhood of their reference IIS gene in the *D. melanogaster* genome assembly (Aug. 2014; BDGP Release 6 + ISO1 MT/dm6). Students then retrieve the protein sequence for the *D. melanogaster* reference gene for a given isoform and run it using *tblastn* against their target *Drosophila* species genome assembly on the NCBI BLAST server (https://blast.ncbi.nlm.nih.gov/Blast.cgi; Altschul et al., 1990) to identify potential orthologs. To validate the potential ortholog, students compare the local genomic neighborhood of their potential ortholog with the genomic neighborhood of their reference gene in *D. melanogaster*. This local synteny analysis includes at minimum the two upstream and downstream genes relative to their putative ortholog. They also explore other sets of genomic evidence using multiple alignment tracks in the Genome Browser, including BLAT alignments of RefSeq Genes, Spaln alignment of *D. melanogaster* proteins, multiple gene prediction tracks (e.g., GeMoMa, Geneid, Augustus), and modENCODE RNA-Seq from the target species. Detailed explanation of how these lines of genomic evidenced are leveraged by students in gene model development are described in Rele et al. (2023). Genomic structure information (e.g., CDSs, intron-exon number and boundaries, number of isoforms) for the *D. melanogaster* reference gene is retrieved through the Gene Record Finder (https://gander.wustl.edu/~wilson/dmelgenerecord/index.html; Rele et al., 2023). Approximate splice sites within the target gene are determined using *tblastn* using the CDSs from the *D. melanogaste*r reference gene. Coordinates of CDSs are then refined by examining aligned modENCODE RNA-Seq data, and by applying paradigms of molecular biology such as identifying canonical splice site sequences and ensuring the maintenance of an open reading frame across hypothesized splice sites. Students then confirm the biological validity of their target gene model using the Gene Model Checker (https://gander.wustl.edu/~wilson/dmelgenerecord/index.html; Rele et al., 2023), which compares the structure and translated sequence from their hypothesized target gene model against the *D. melanogaster* reference gene model. At least two independent models for a gene are generated by students under mentorship of their faculty course instructors. Those models are then reconciled by a third independent researcher mentored by the project leaders to produce the final model. Note: comparison of 5’ and 3’ UTR sequence information is not included in this GEP CURE protocol.” (Gruys et al, 2025)

## Supporting information

FASTA, GFF, and PEP Files

## Supplemental Material

1. Zip file containing FASTA, PEP, GFF files for the gene model
2. Figure 1 in high resolution

## Metadata

Bioinformatics, Genomics, *Drosophila*, Genotype Data, New Finding

## Acknowledgements

We thank Wilson Leung (Washington University, St. Louis) for developing and maintaining the technological infrastructure that supported the creation of this gene model, as well as Chinmay Rele and Laura K. Reed (University of Alabama) for their guidance and encouragement throughout the project. We are also grateful to FlyBase for providing the authoritative database for *Drosophila melanogaster* gene models. FlyBase is supported by grants NHGRI U41HG000739 and U24HG010859, UK Medical Research Council MR/W024233/1, NSF 2035515 and 2039324, BBSRC BB/T014008/1, and Wellcome Trust PLM13398.

